# Regional transmission and reassortment of 2.3.4.4b highly pathogenic avian influenza (HPAI) viruses in Bulgarian poultry 2017/18

**DOI:** 10.1101/2020.04.14.040592

**Authors:** Divya Venkatesh, Adam Brouwer, Richard Ellis, Gabriela Goujgoulova, James Seekings, Ian H. Brown, Nicola S. Lewis

## Abstract

Between 2017 and 2018, several farms across Bulgaria reported outbreaks of H5 HPAI viruses. In this study we use genomic and traditional epidemiological analyses to trace the origin and subsequent spread of these outbreaks within Bulgaria. Both methods indicate two separate incursions, one restricted to the North-Eastern region of Dobrich, and another largely restricted to Central and Eastern Bulgaria including places such as Plovdiv, Sliven and Stara Zagora, as well as one virus from the Western region of Vidin. Both outbreaks likely originate from different European 2.3.4.4b virus ancestors circulating in 2017. The viruses were likely introduced by wild birds or poultry trade links in 2017 and have continued to circulate, but due to lack of contemporaneous sampling and sequences from wild bird viruses in Bulgaria, the precise route and timing of introduction cannot be determined. Analysis of whole genomes indicates complete lack of reassortment in all segments but the MP, which presents as multiple smaller clusters associated with different European 2.3.4.4b viruses. Ancestral reconstruction of host states of the HA gene of viruses involved in the outbreaks suggests that transmission is driven by domestic ducks into galliform poultry. Thus, according to present evidence we suggest that surveillance of domestic ducks as epidemiologically relevant species for subclinical infection. Monitoring spread due to movement between farms within regions and links to poultry production systems in European countries can help to predict and prevent future outbreaks.

## Introduction

Aquatic birds form the reservoir for avian influenza viruses, where multiple subtypes circulate and generally do not cause any disease. Periodically, such viruses can infect gallinaceous poultry as low pathogenic avian influenza (LPAI) viruses. Here, viruses of H5 and H7 HA subtypes can mutate into highly pathogenic forms (Bosch *et al*., 1979; Röhm *et al*., 1995; Alexander, 2000; Alexander, 2007). Highly pathogenic avian influenza (HPAI) viruses constitute a major threat to poultry populations worldwide because of their high levels of mortality, potential for spread, impact on livestock production, and their zoonotic potential.

Such LPAI to HPAI transformations have occurred on multiple occasions via introduction of multiple basic amino acids in the hemagglutinin (HA) proteolytic cleavage site. After influenza viruses enter the cell, there is cleavage of the inactive precursor HA0 protein at this site, into HA1 and HA2 subunits. The regular cleavage site is cleaved by trypsin or trypsin-like proteases present largely in the respiratory and intestinal epithelia. However, the multi-basic cleavage site can be cleaved by several ubiquitous proteases such that the virus can infect and replicate in multiple tissue types resulting in a systemic disease with high morbidity and mortality (Klenk and Rott, 1988; Rott, 1992).

Outbreaks caused by certain HPAIs of the A/goose/Guangdong/1/1996 (Gs/GD) lineage of H5 viruses continued to circulate in domestic birds in some countries and were re-introduced into wild aquatic bird reservoirs to spread to new geographic areas (Sonnberg *et al*., 2013; Lycett *et al*., 2019). Thus far, viruses bearing this Gs/GD lineage HA have caused infections in poultry, wild birds and humans in up to 83 countries across several continents (EMPRES-i data http://empres-i.fao.org/eipws3g/ accessed 9 April 2019, (Claes *et al*., 2014)). Continued circulation of viruses bearing this H5 HA gene in poultry and wild aquatic reservoirs has resulted in its diversification into multiple clades and sub-clades; by 2012, 12 distinct clades were identified (World Health Organization/World Organisation for Animal Health/Food and Agriculture Organization (WHO/OIE/FAO) H5N1 Evolution Working Group, 2014). Viruses bearing HAs of the Gs/GD 2.3.4 lineage emerged around 2009 – 2013 revealed an early propensity to reassort with NA subtypes other than N1, unlike earlier clades, and showed unprecedented geographical range expansion via poultry trade and wild bird migration (Claes *et al*., 2016; Dhingra *et al*., 2016; Lee *et al*., 2017). Since 2014, HPAI clade 2.3.4.4 viruses have spread rapidly through Eurasia and into North America via migratory wild aquatic birds and have evolved through reassortment with prevailing local low pathogenicity avian influenza viruses. They are associated with variable disease severity, including subclinical infection in wild birds and domestic waterfowl (European Food Safety Authority, 2014). From May 2016, the clade 2.3.4.4 Group B (2.3.4.4b) H5N8 viruses re-emerged in Europe causing numerous outbreaks in poultry and massive die-offs in wild birds (Beerens *et al*., 2017; Kleyheeg *et al*., 2017; Pohlmann *et al*., 2017; Poen *et al*., 2019). Several reassortment events led to the emergence and detection of HPAI H5N5 in several European countries, Georgia and Israel between November 2016 and June 2017 (Alarcon *et al*., 2018) and HPAI H5N6 in Greece in February 2017 (OIE, 2017), with the subsequent wave of HPAI H5N6 viruses evolving from the H5N8 2016–17 viruses during 2017 by reassortment of a European HPAI H5N8 virus and wild host reservoir LPAI viruses (Kwon *et al*., 2018; Poen *et al*., 2019).

With over 1,197 H5 HPAI outbreaks reported in poultry or captive birds in 20 countries, the 2016/2017 HPAI epidemic was the largest ever recorded in the EU in terms of number of outbreaks, geographic distribution and the number of dead wild birds (Brown *et al*., 2017b; Alarcon *et al*., 2018). Reports indicate that the H5N8 virus persisted during early winter 2016 into the late summer 2017 at least, leading to in sporadic outbreaks in poultry and wild bird infections towards autumn (Brown *et al*., 2017b). Events decreased in early winter 2017 with 48 poultry outbreaks and 9 wild bird detections recorded towards the end of 2017 in winter, with even fewer outbreaks and detections in wild birds the following year (Brown *et al*., 2017b; Brown *et al*., 2017a).

Viruses of H5 subtype were largely absent through 2018 and past February until December 2019 only, two wild bird HPAI events were identified both in Denmark, but Bulgaria remained the only country in Europe reporting HPAI outbreaks in poultry holdings (Adlhoch *et al*., 2018; Adlhoch *et al*., 2019). In this study we use genomic and traditional epidemiological methods to source and track the spread of HPAI poultry outbreaks that occurred in Bulgaria during 2017/18. We aim to understand the role of wild birds and the poultry system structure in HPAI introduction, onward spread in poultry and the subsequent potential for endemic circulation of highly pathogenic viruses in domestic birds to help design better containment and preventive measures.

## Methods

### Epidemiology

Epidemiological data on infected premises were obtained from the Animal Disease Notification System managed by the European Commission and validated with the National Diagnostic Research Veterinary Medical Institute in Bulgaria. Isolates obtained from the veterinary authorities in Bulgaria were matched to the appropriate infected premise and GIS analyses was performed using ArcGIS Desktop 10.2.2 (Environmental Systems Research Institute, 2014).

Additional background data were also taken from Bulgarian presentations to the Standing Committee on Plants, Animals, Food and Feed (PAFF, 2018).

### Sequencing

RNA was sequenced using the MiSeq platform. Briefly, viral RNA was extracted from the nasal swab using the QIAmp viral RNA mini kit without the addition of carrier RNA (Qiagen, Manchester, UK). cDNA was synthesised from RNA using a random hexamer primer mix and cDNA Synthesis System (Roche, UK). The Sequence library was prepared using a NexteraXT kit (Illumina, Cambridge, UK). Quality control and quantification of the cDNA and Sequence Library was performed using Quantifluor dsDNA System (Promega, UK). Sequence libraries were run on a Miseq using MiSeq V2 300 cycle kit (Illumina, Cambridge, UK) with 2 × 150 base paired end reads. The raw sequence reads were analysed using publicly available bioinformatics software, following an in-house pipeline, available on github (https://github.com/ellisrichardj/FluSeqID/blob/master/FluSeqID.sh). This pipeline de novo assembles the raw data using the Velvet assembler (Zerbino and Birney, 2008), Basic Local Alignment Search Tools (BLASTs) the resulting contigs against a local database of influenza genes using Blast+ (Camacho *et al*., 2009),then maps the raw data against the highest scoring blast hit using the Burrows-Wheeler Aligner (Li, 2013). The consensus sequence was extracted from the resultant bam file using a modified SAMtools software package (Li *et al*., 2009), script (vcf2consensus.pl) available at: https://github.com/ellisrichardj/csu_scripts/blob/master/vcf2consensus.pl. Whole-genome sequences of the viruses are available on GISAID database with unique IDs:

EPI_ISL_419212, EPI_ISL_419344, EPI_ISL_419345, EPI_ISL_419346,

EPI_ISL_419347, EPI_ISL_419348, EPI_ISL_419349, EPI_ISL_419350,

EPI_ISL_419351, EPI_ISL_419352, EPI_ISL_419353, EPI_ISL_419354,

EPI_ISL_419355, EPI_ISL_419356, EPI_ISL_419357, EPI_ISL_419358,

EPI_ISL_419359, EPI_ISL_419360, EPI_ISL_419361, EPI_ISL_419362,

EPI_ISL_419363, EPI_ISL_419364, EPI_ISL_419365, EPI_ISL_419363,

EPI_ISL_419367,

### Visualising reassortment

We first used BLAST on GISAID (Elbe and Buckland-Merrett, 2017; Shu and McCauley, 2017) to identify the closest genetic relatives to the Bulgarian outbreak viruses. We found that for all segments, the top 50 blast hits were largely from viruses bearing H5 HA and isolated from avian hosts in Europe in the year 2016 or later. Therefore, for the reassortment analysis we downloaded whole-genome sequences from all H5Nx viruses isolated between January 2016 and December 2018 from avian hosts in Europe from FluDB and GISAID. Sequences from each segment were checked for quality including sequence length > 30% average length for a given segment, removing duplicates, and all 8 segments present. The resulting datasets were combined with Bulgarian segment sequences.

Final datasets were aligned using MAFFT v7.305b (Katoh and Standley, 2013) and trimmed to only retain nucleotides from the starting ATG until the final STOP codon. We inferred Maximum Likelihood (ML) phylogenetic trees for each gene segment using IQ-TREE, 1.5.5 (Nguyen *et al*., 2015) and obtained branch supports with Shimodaira-Hasegawa (SH)-like approximate Likelihood Ratio Test (aLRT, 1,000 replicates).

BALTIC (backronymed adaptable lightweight tree import code, https://github.com/evogytis/baltic) was used to compare the phylogenetic structure of the segment genes. The phylogenetic position of each strain was traced, coloured according to the lineage and location across unrooted ML trees for HA and all internal gene segments. Figures were generated by modifying scripts from a similar analysis (Bell and Bedford, 2017) and editing in Adobe Illustrator. We selected a qualitative palette of colours using http://colorbrewer2.org/.

### Spatial phylodynamic analysis and ancestral host reconstruction

For the H5 analysis, 1189 HA sequences of strains isolated between January 2016 and December 2018 were downloaded from GISAID (Supplemental Table 1).

The individual H5 dataset was first subjected to a quality control step where all duplicate sequences and sequences bearing duplicate IDs were removed (where 871 sequences remained). These sequences together with those sequenced from Bulgaria (BGR) were aligned using MAFFT v7.305b (Katoh and Standley, 2013) and used in the FastTree program (Price *et al*., 2009) to generate maximum-likelihood tree with GTR+gamma substitution model. All sequences from taxa outside of the fully supported (100%) cluster with BGR were discarded. The H5 HA dataset included representatives from the HPAI clade 2.3.4.4b (H5N8) 2016-17 clade (need WHO H5 group nomenclature reference) to form a new dataset of 104 sequences from which a final H5 ML tree was inferred using FastTree. This tree was analysed with tempest v1.5 to check for clock-like behaviour (Rambaut *et al*., 2016).

Bayesian phylogenetic trees were inferred using BEAST v1.10.4 (Suchard *et al*., 2018) to determine the time of emergence of the Bulgarian viruses. We coded each taxon with host (anseriformes, galliformes, other) and location (Dobrich, Haskovo, Plovdiv, Sliven, Stara Zagora, Yambol) information. AICM (Akaike’s information criterion through Markov chain Monte Carlo) values were used to select the appropriate state (symmetric vs asymmetric) and clock (strict vs relaxed (uncorrelated relaxed lognormal)) models. Asymmetric state transition and strict clock were chosen over symmetric/relaxed clock. GMRF Bayesian Skyride population prior was used with a random starting tree. All other priors were set to default. MCMC was set to 50,000,000 generations. Two separate runs were performed to ensure convergence between runs. Log files were analysed in Tracer v1.7.1 to determine convergence, and to check that ESS values were beyond threshold (>200). Log and trees files from both runs were combined using Log Combiner v 1.10.4. Tree annotator v1.10.4 was used to generate a maximum credibility tree (MCC) using 10% burn in and median node heights. The MCC tree was then annotated to include posterior probability values and time scales, and host and location states, and plotted in R v 3.5 using the ggtree package (Yu *et al*., 2017).

## Results

### Epidemiology

Between 17/10/2017 and 08/04/2019, Bulgaria reported 38 HPAI H5 outbreaks in poultry. Bulgaria accounted for 43.6% of outbreaks in Europe in this timeframe. During this period there were no positive wild birds detected in Bulgaria among the 47 and 58 dead or moribund wild birds sampled through passive surveillance in 2017 and 2018 respectively (Brouwer *et al*., 2019). We observed four loose geographical clusters in the infected premise locations; Plovdiv and Haskovo in Central Bulgaria, Yambol in East Bulgaria and Dobrich in North-East Bulgaria and sporadic cases in Vidin and Lovech in North-West Bulgaria. A range of species were affected including ducks (19), chickens (7), turkeys (1) partridges (1), and backyard poultry including hens and ducks (10). There was no specific-species bias between clusters. Both backyard and commercial settings were affected. Of the 38 infected premises, samples from 25 locations were sent to APHA for confirmatory testing, virus isolation isolation and genetic analyses. Samples from infected premises detected after 31/10/2018 were not sent for confirmatory testing, and APHA did not receive samples from 2 outbreaks in ducks confirmed on the 10/4/2018 and 24/5/2018 in Plovdiv (Submission ID 2018/3 and 2018/8).

**Figure 1:**
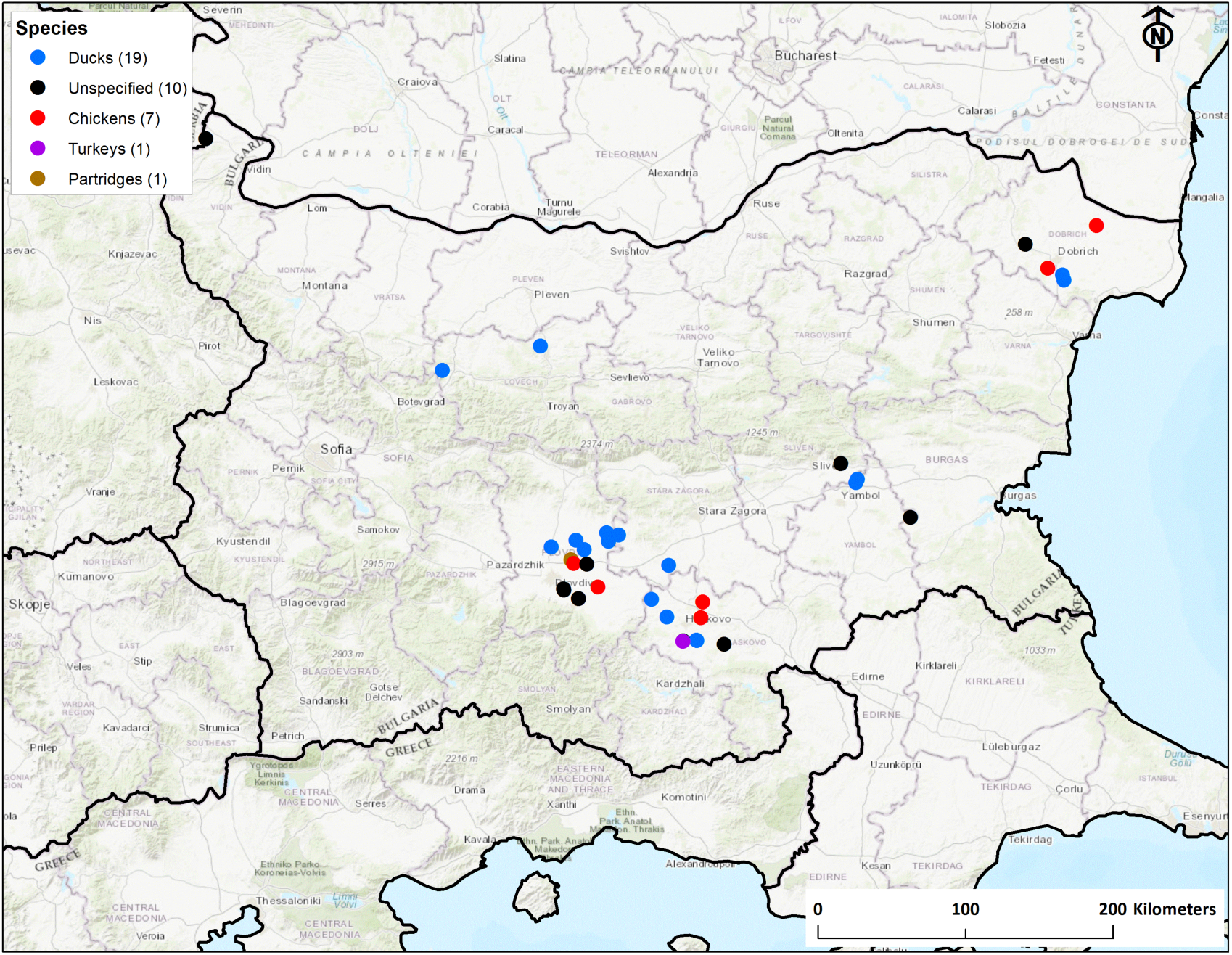
H5N8 HPAI outbreaks in poultry in Bulgaria 2017-2019. Colour of dots indicates species affected as shown in the key. Numbers in brackets indicate the number of premises in which each species was affected.

Clinical indicators especially including mortality typical of HPAI were sufficient triggers for vet investigation. Mortality in Anseriformes birds was more mixed with a number of early events showing mortality, but later events showing little or no mortality. As **Figure 2** shows, commercial duck production in Bulgaria is concentrated in the South/Central and East parts of the country with the North and West having relatively few registered holdings, whilst chicken holdings are more homogenously located throughout the country. Virologically-positive poultry-infected premises were found primarily in the high duck density areas of Bulgaria, for both duck outbreaks as well as gallinaceous poultry outbreaks.

### Poultry and wild bird surveillance

Surprisingly, results from the EU active serological surveillance programme in Bulgaria for poultry show no positive serological results between 2016 and 2018, despite sampling around 500 poultry holdings a year of which 109, 276 and 155 duck holdings were sampled respectively in 2016, 2017 and 2018. As part of their outbreak response, Bulgarian authorities conducted an additional reactive serological programme in all farms with domestic poultry, which sampled broilers at 2 monthly intervals and other poultry at age 50-60 days. Of those sampled, there were 31 serology positive holdings and one flock was also shown to be actively infected using PCR.

**Figure 1A.**
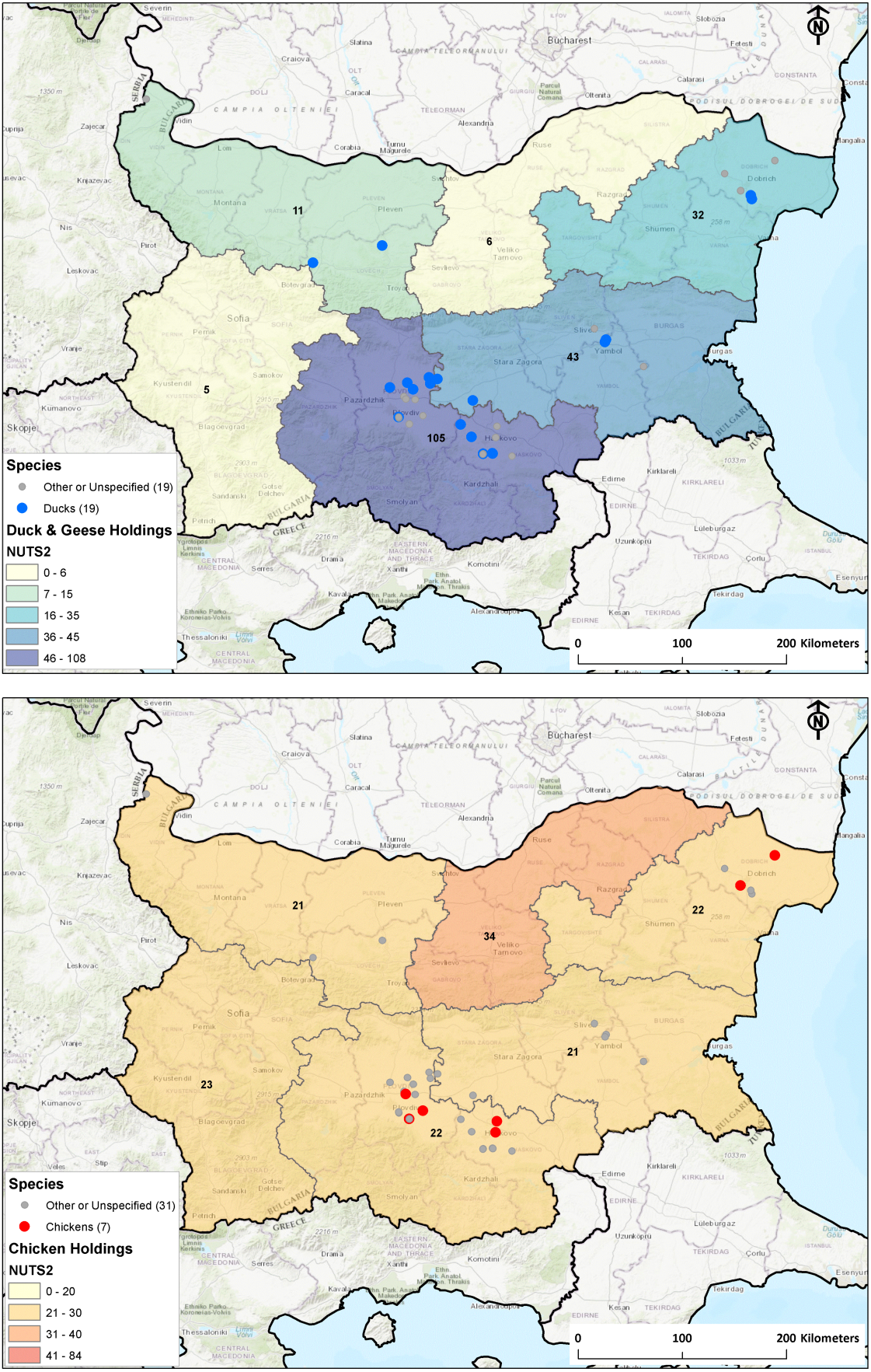
(top) and 2B (bottom): 2017-2019 Poultry outbreaks in Bulgaria. Highlighted areas are NUTS2 areas with labels indicating the number of Duck and Geese registered holding (top, blue) and number of Chicken registered holdings (bottom, red)

There was also a compulsory programme of passive surveillance in dead or moribund wild birds that is co-financed by the EU (as stated by European Union guidelines (EC, 2010)). Results from this surveillance for 2017 and 2018 are shown in **Table 1** alongside results from the two EU member-states of Romania and Greece that share a land border with Bulgaria. Whilst there were initial detections of H5N8 HPAI in south-east Europe in late 2016, these continued and accelerated in early 2017 with the last confirmed H5N8 case in wild birds found in Romania in March 2017. There were no identified cases of H5N8 found in wild birds in the region throughout the remainder of 2017 and no cases identified in 2018.

**Table 1:** Wild Bird Surveillance results from Bulgaria, Romania and Greece (2017-2018)

|  | 2016 |  |  | 2017 |  |  | 2018 |  |  |
| --- | --- | --- | --- | --- | --- | --- | --- | --- | --- |
| Country | Number of birds sampled | Total number of HPAI H5 detections | Total number of HPAI H5N8 detections | Number of birds sampled | Total number of HPAI H5 detections | Total number of HPAI H5N8 detections | Number of birds sampled | Total number of HPAI H5 detections | Total number of HPAI H5N8 detections |
| Bulgaria | 9 | 2 | 2 | 47 | 14 | 14 | 58 | 0 | 0 |
| Greece | 16 | 1 | 1 | 90 | 13 | 12 | 13 | 0 | 0 |
| Romania | 275 | 10 | 10 | 528 | 162 | 143 | 244 | 0 | 0 |

**Figure 3:**
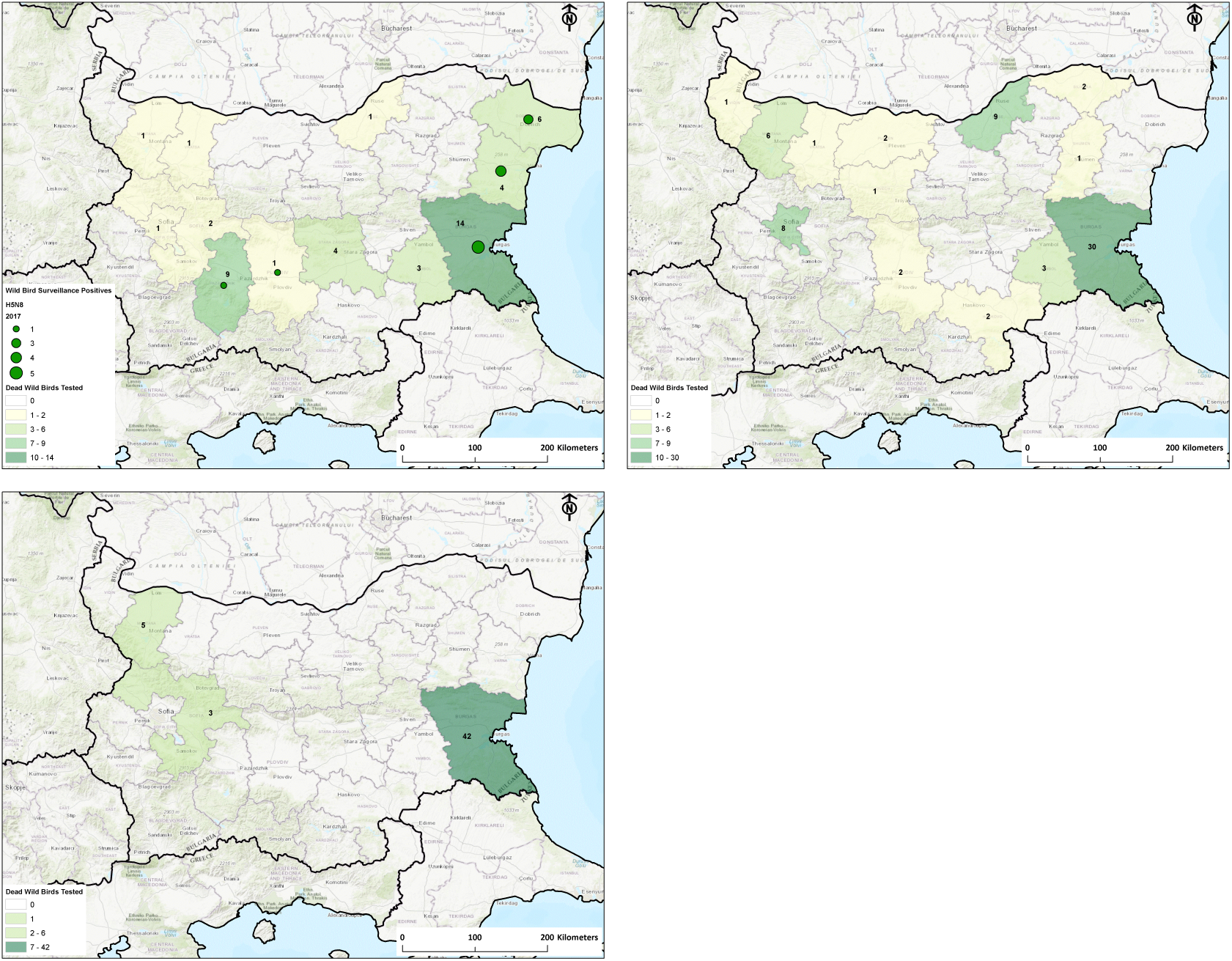
Clockwise from Top Left: Surveillance of dead wild birds in Bulgaria from 2017, 2018 and 2019 (first six months). Green circles represent H5N8 wild bird positives, regional shading indicates number of dead wild birds tested.

### Genetic structure of Bulgarian H5 HA

The maximum-likelihood tree in **Figure 4** shows that the highly pathogenic H5 HAs from Bulgaria arose from European H5N8 strains of the 2.3.4.4b lineage. They show no close genetic link with the subsequent H5N6 wave from Asia that spread into Europe (Kwon *et al*., 2018; Poen *et al*., 2019). Bulgarian virus HAs isolated in 2017-18 form two separate clusters originating from distinct 2.3.4.4b H5 virus ancestors, together with a lone 2017 strain from Dobrich. The clusters are structured geographically with strains exclusively from Dobrich forming one cluster in the North East (in green). Strains from Central Bulgarian regions Plovdiv, Haskovo, and Stara Zagora (in red), Eastern regions Yambol and Sliven (in blue), and Vidin in the West (in purple) form a single mixed cluster. This implies that viruses in both clusters are circulating separately and their HAs likely have separate origins. Within both clusters, sub-clusters with possible separate origins are discernible but the data is likely confounded by under-sampling.

**Figure 4:**
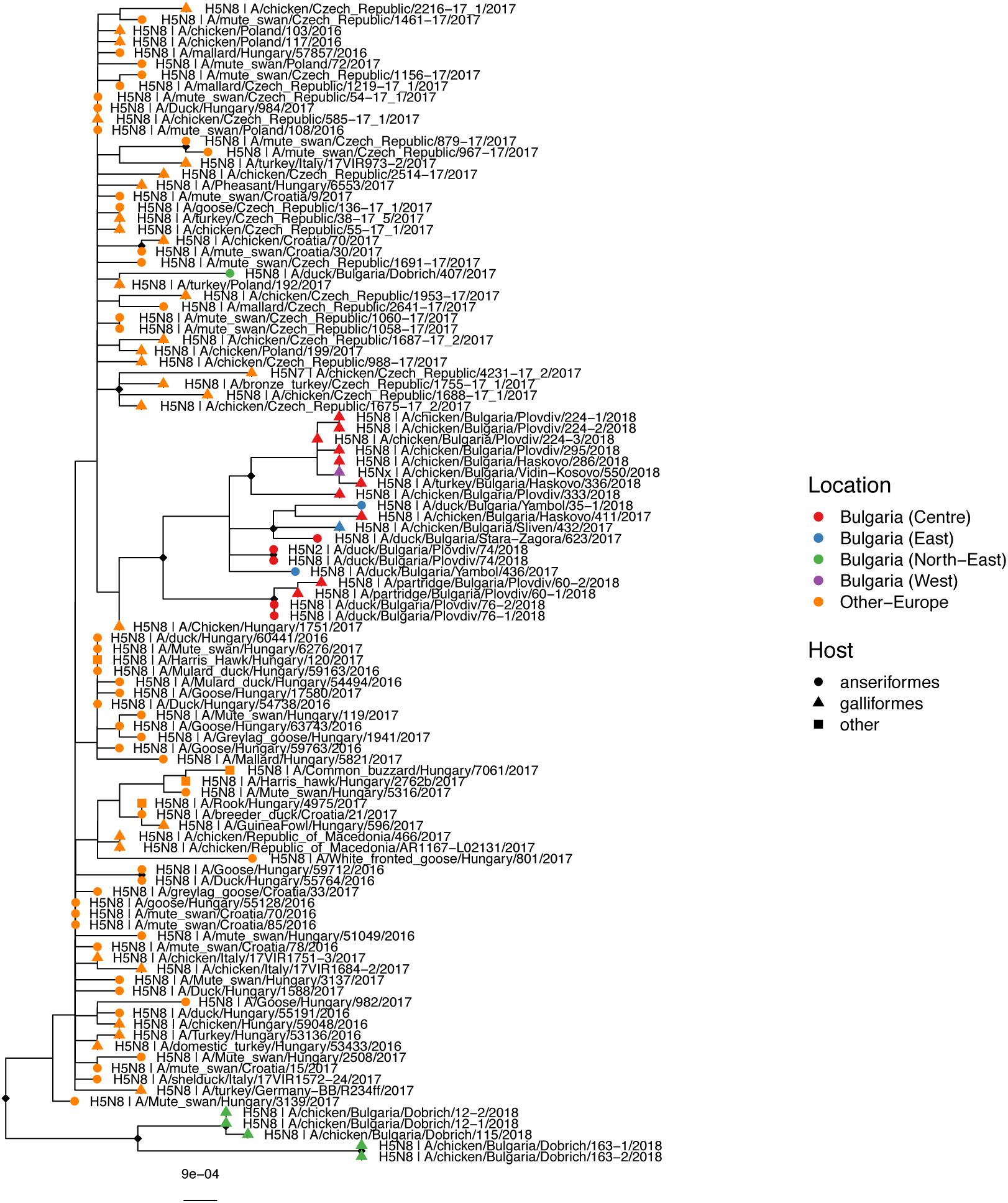
Mid-point rooted maximum-likelihood tree showing phylogenetic relationships between HA sequences from viruses isolated from poultry outbreaks in Bulgaria (2017-18). Tip shapes indicate host, while colours indicate location from which the virus was sampled. Diamond shapes at the nodes branch support values > 90/100.

### Timing of introductions, host dynamics and spatial spread

The BEAST maximum clade credibility (MCC) tree for the HA gene **(Figure 5A)** shows that after introduction, virus HAs in the mixed cluster have been circulating within Bulgaria since approximately early 2017 (95% HPD: September 2016 – May 2017) and those in the Dobrich cluster, since May 2017 (95% HPD: March – October 2017). This means that after the two separate introductions into Bulgarian poultry, the viruses have been transmitted locally within the country. The lone October 2017 Dobrich virus HA was a separate introduction which failed to spread.

**Figure 5A:**
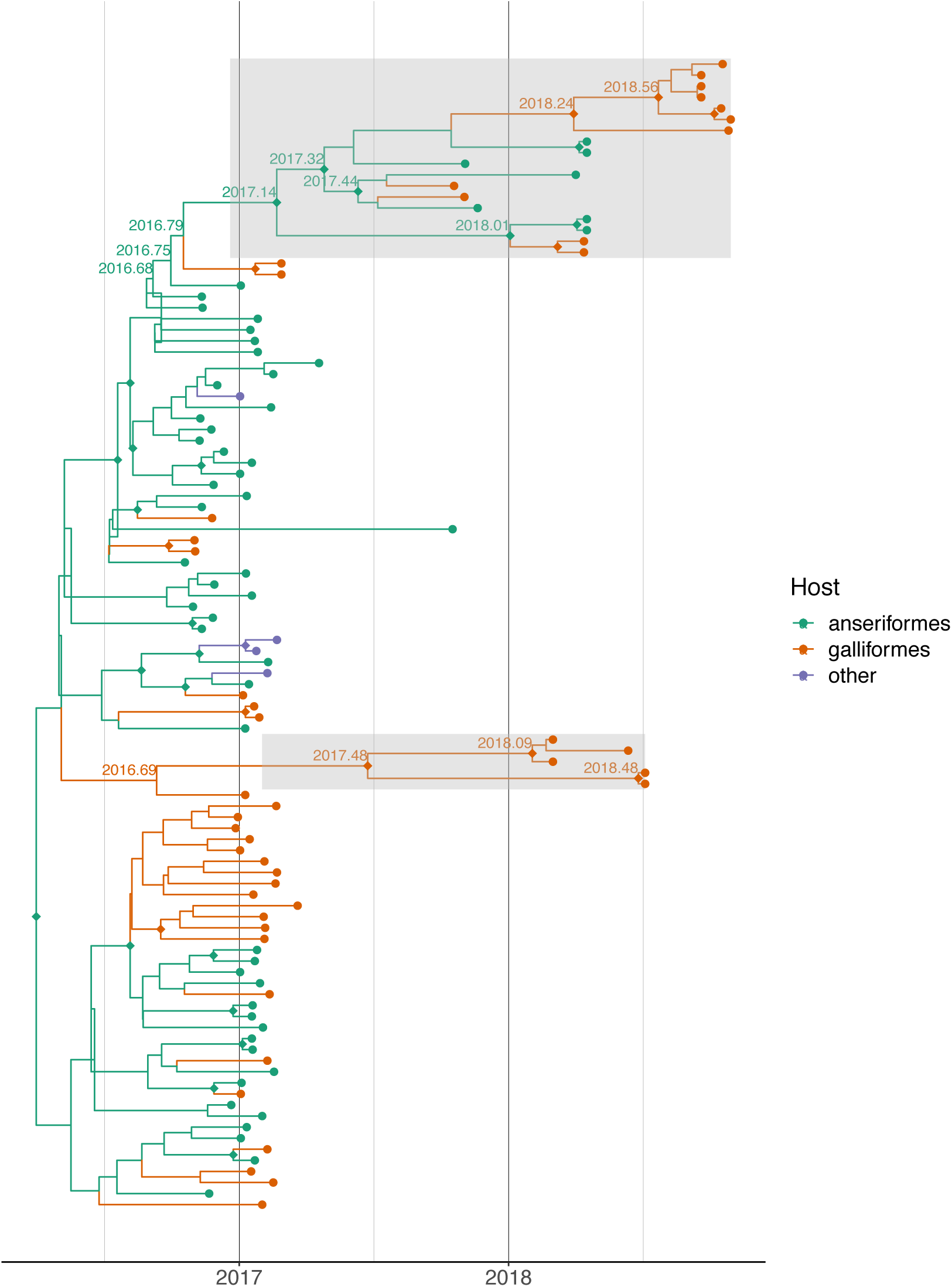
BEAST maximum clade credibility tree showing reconstruction of host states at putative ancestral nodes of viruses isolated from poultry outbreaks in Bulgaria (2017-18, clusters highlighted gray). Tip and branch colours indicate host states. Selected nodes of interest are labelled with putative times to most recent common ancestor (TMRCAs). Diamond shapes at the nodes indicate posterior probability values > 85/100.

Location-wise, it is likely that the source of the mixed cluster of viruses was Plovdiv, from where it spread to other Central and Eastern Bulgarian regions **(Figure 5B and 6)**. While the monophyly of the Bulgarian mixed and Dobrich clades are very well-supported (posterior probability (pp) = 0.999), the identity of closest related virus is unclear for both the mixed and Dobrich clusters (pp = 0.046, 0.478) likely due to lack of sampling. Even though 2 and 14 HPAI H5 viruses were detected in wild birds in Bulgaria in 2016 and 2017 respectively, we do not have sequences from these viruses to determine how they relate to the poultry outbreaks. No onward transmission to other geographic areas were detected as of early 2019. Although there is evidence of serologically positive poultry within Bulgaria in 2018-19, we were unable to explore their provenance due to lack of viruses or genetic sequences.

**Figure 5B:**
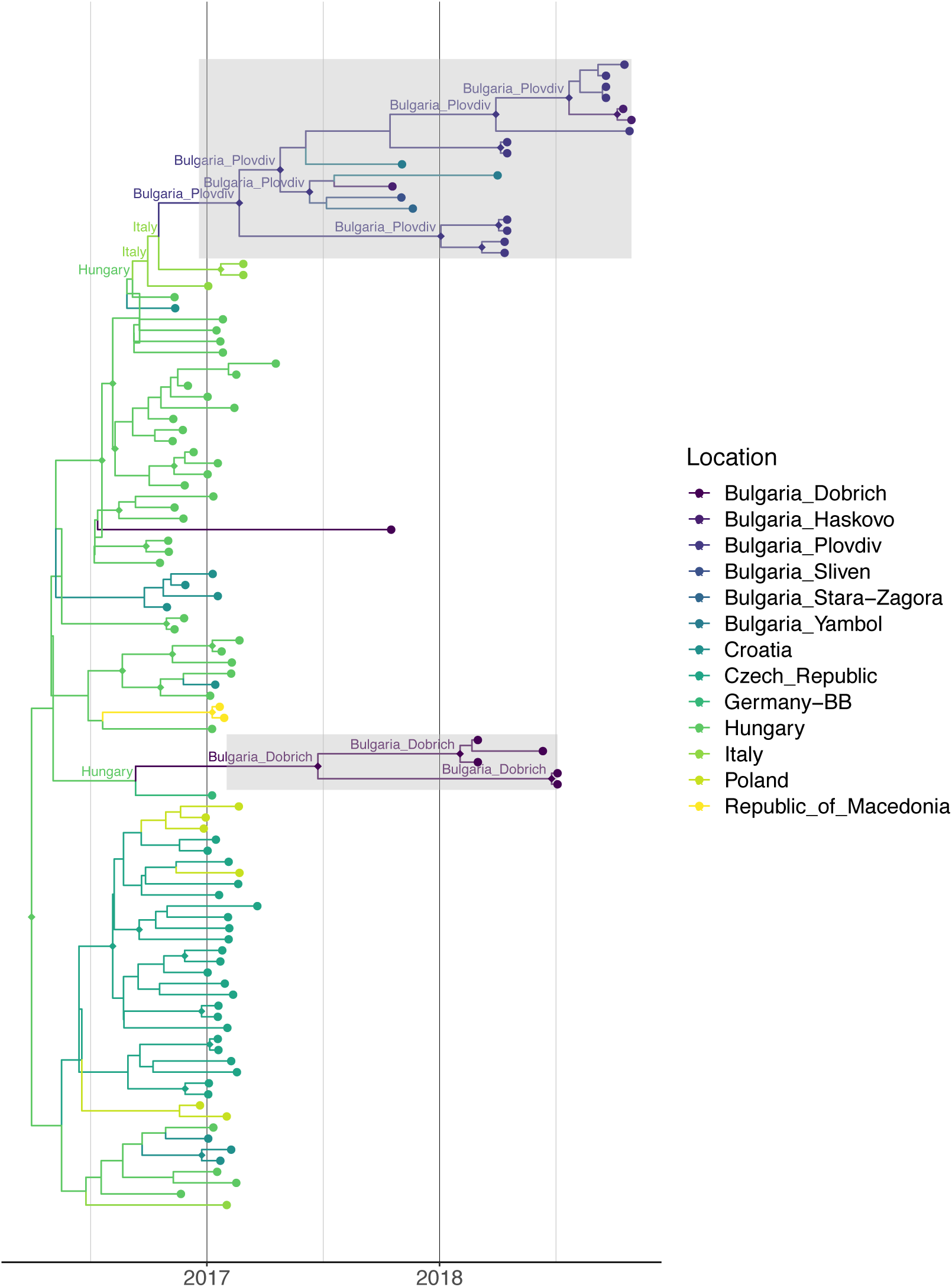
BEAST maximum clade credibility tree showing reconstruction of location states at putative ancestral nodes of viruses isolated from poultry outbreaks in Bulgarian (2017-19, highlighted clusters in gray). Tip and branch colours indicate location states. Selected nodes of interest are labelled with putative location states. Diamond shapes at the nodes indicate posterior probability values > 85/100.

**Figure 6:**
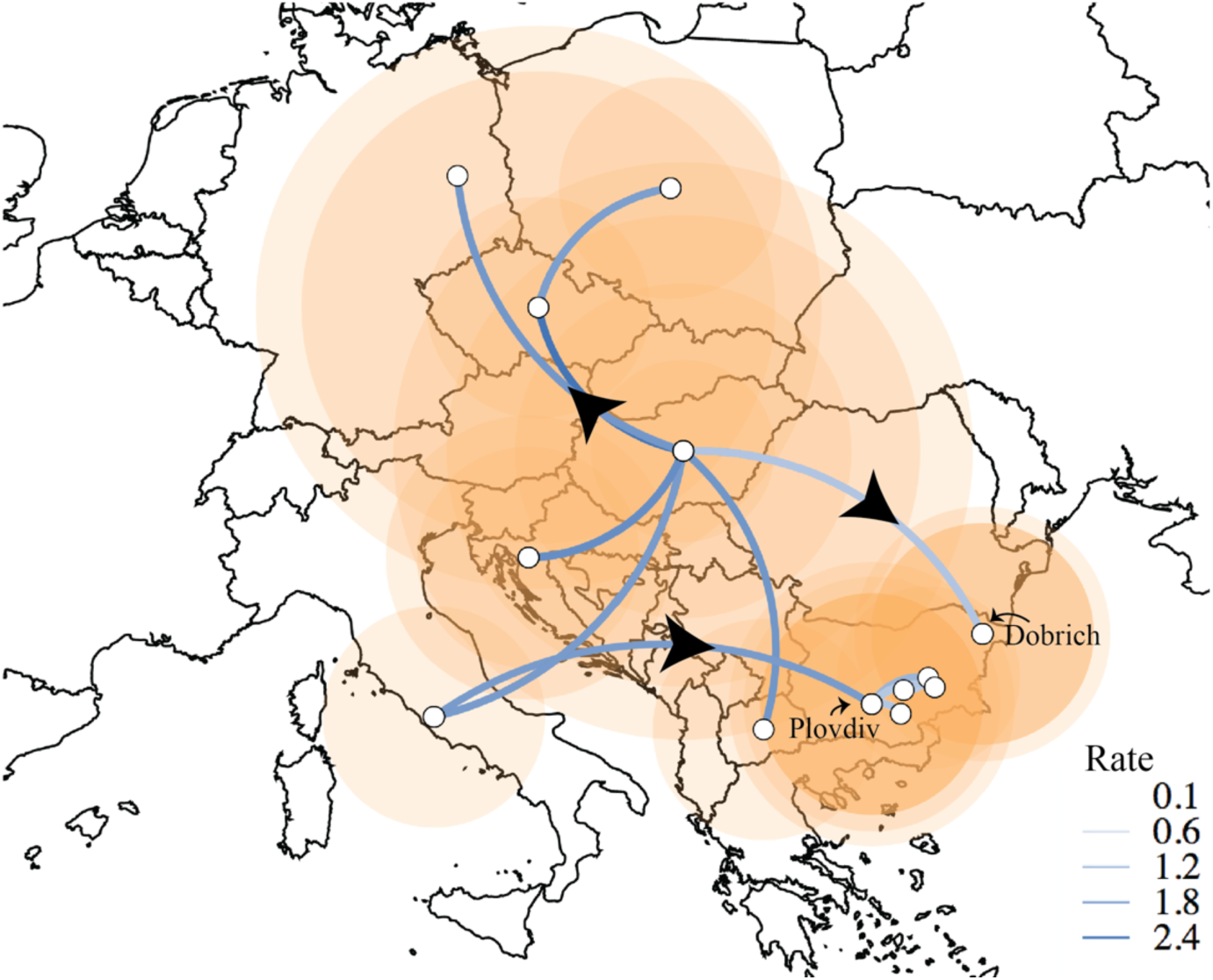
Summary of transmission rates between different locations as calculated from BEAST analysis using SpreaD3. Arrows show direction of transmission, size of orange circles corresponds to cumulative number of cases.

Within poultry, the transmission occurs largely in the direction from ducks (anseriformes) to chicken (galliformes), but not from chickens back into ducks **(Figure 7)**. This pattern is consistent with our epidemiological findings above as well as previous studies in wild birds (Venkatesh *et al*., 2018) and other reported 2.3.4.4b poultry outbreaks in Europe which implicated ducks in local spread (Marinova-Petkova *et al*., 2016; Guinat *et al*., 2020).

**Figure 7:**
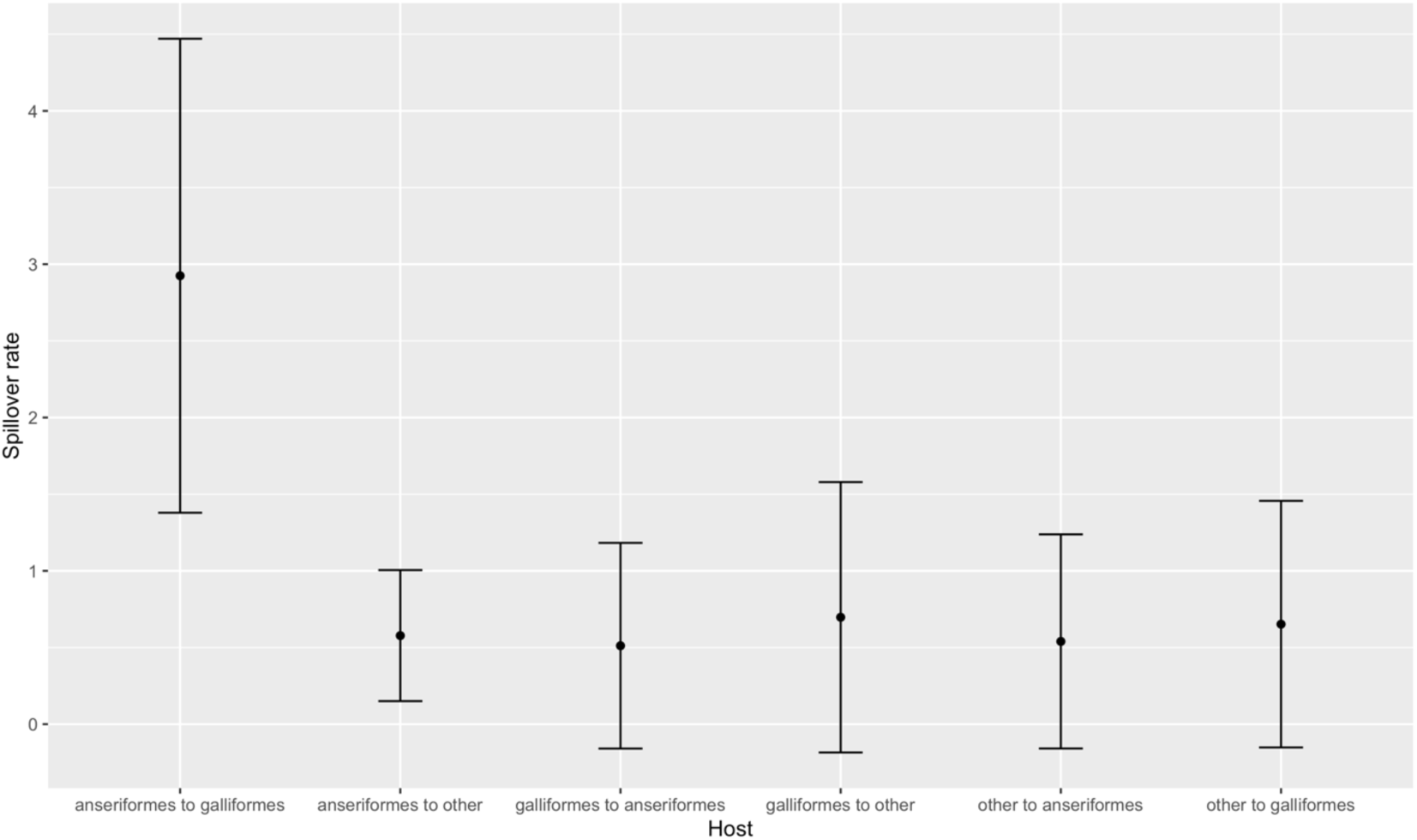
Summary of spill over rates between different hosts (Anseriformes, Galliformes, other) as calculated from BEAST analysis. Error bars indicate 95% confidence intervals.

### Whole-genome analysis

We used BALTIC (backronymed adaptable lightweight tree import code, https://github.com/evogytis/baltic) to compare the phylogenetic structure of the internal genes of the Bulgarian strains compared to other HPAI H5 and LPAI viruses. To visualise incongruence, the phylogenetic position of each sequence (coloured according to the origin of its HA) was traced across all eight trees (**Figure 8**). Parallel lines indicate similar origin, whereas crossed lines indicate differential ancestry. We find that the internal genes of the Bulgarian strains are largely derived from the same ancestral viruses, the lines connecting the strains are largely parallel and remain within the same larger cluster of H5 viruses. Some variation is present for the MP gene, which shows multiple clusters with each related to MP segments associated with other European 2.3.4.4b viruses isolated from hosts such as duck, mute swan and pheasant (Supplementary Figure 2) – if there are any Bulgarian intermediaries, they remain unsampled. Like the HAs, none of the segments appear to be related to any viruses from the recent 2.3.4.4b H5N6 wave in Asia that spread into Europe, but to the dominant 2.3.4.4b H5N8 lineage.

**Figure 8:**
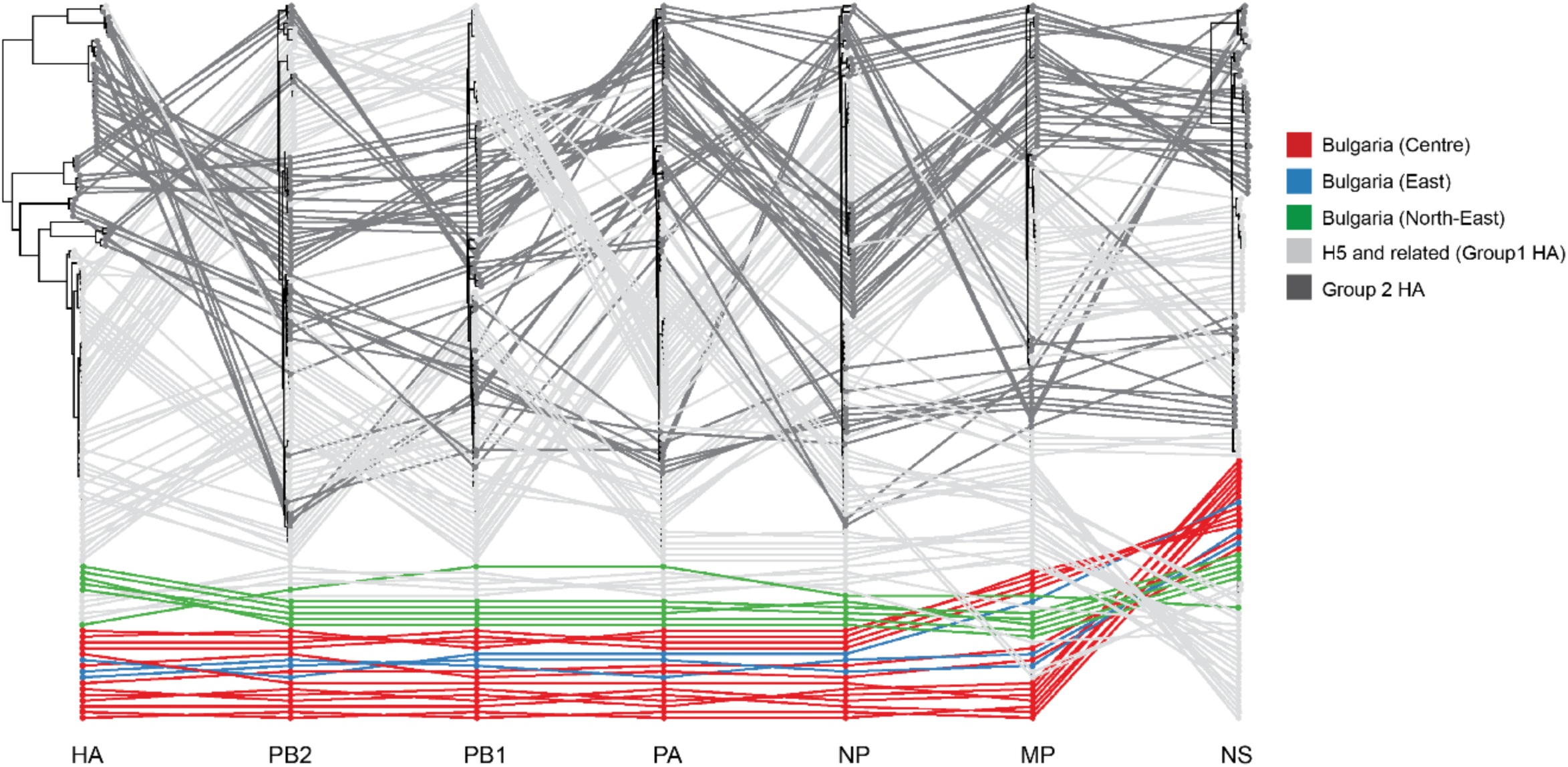
Visualisation of phylogenetic incongruence as an indicator of differential ancestry of segment sequences i.e., reassortment. Maximum-likelihood trees for each segment are plotted with lines connecting the same virus strain across all trees. Colours indicate origin of viruses sampled in Bulgaria (red, blue, green, purple) during 2017-18 or related sequences outside of Bulgaria (grey).

This pattern of limited reassortment is consistent with virus transmission and endemic maintenance within Bulgarian poultry and no epidemiologic role for currently circulating low pathogenic avian influenza in wild birds.

## Discussion

In this study we investigated an apparently unique and independent epidemiological event in Bulgaria in 2017-18. During this study period there was no evidence of sustained transmission or endemicity of other H5N8 2.3.4.4b virus outbreaks in Europe during 2018-19. Furthermore, we were able to trace the Bulgarian outbreaks during 2017-18 to the 2.3.4.4b lineage viruses circulating in Europe 2017. As in a previous Bulgarian study (Marinova-Petkova *et al*., 2016) we found no links to viruses from Bulgarian wild birds despite concurrent sampling in the earlier study; nor do we find links to Eurasian/East Asian wild birds. We demonstrated incursion of these viruses into Bulgarian domestic birds has occurred on at least two occasions into different regions of the country, from distinguishable genetically distinct 2.3.4.4b ancestral strains which, after introduction, have circulated with spatial separation within country since 2017. Phylogenetically long branch lengths connecting to poultry viruses also indicates significant in-country circulation of undetected disease and unsampled strains for a period before outbreaks. From available data we cannot say if the route of transmission was via migratory wild birds or duck trade links between Bulgaria and other European countries.

During the outbreaks, we show that transmission into galliform poultry is likely driven by domestic ducks. Given the presence of limited reassortment signals (**Figure 8**) and absence of detections in wild birds in Bulgaria or any other parts of Europe during 2018, H5N8 viruses have been likely maintained endemically within the domestic bird sector. However, for the MP gene segment (see **Figure S2**) we did find multiple clusters of viruses, each associated with isolates from multiple species (ducks, swan, pheasant) in different countries (Netherlands, England, Hungary). A wider sampling of domestic ducks might reveal if the limited reassortment that was detected in the MP gene segment is driven by transmission from these hosts. Additionally, H5N8 viruses identified in 2019 in locations such as Poland and Slovakia appear to be unrelated to Bulgarian viruses (OFFLU, 2020). Although we do not have sequences yet, H5 HPAI viruses continue to be detected in Bulgarian poultry (as per information downloaded from http://empres-i.fao.org/eipws3g/, see **Supplementary Table 1**). Once sequences are available, genetic analysis will reveal whether they are related to the viruses we describe in this study, either revealing continued transmission or a new introduction from wild birds.

We note the risk of the domestic duck sector as a source of cryptic spread and maintenance after introduction. Our results are consistent with detailed spatial modelling during the 2016-17 HPAI 2.3.4.4b outbreaks in France (Guinat *et al*., 2019; Guinat *et al*., 2020), and previous surveillance in Bulgaria during 2008-12 (Marinova-Petkova *et al*., 2016). Both studies found a major role for duck production systems, particularly foie gras, in the spread of avian influenza in the poultry sector. France and Bulgaria together with Hungary are the major producers of foie gras in Europe and maintain trade links connecting duck farms (transport of ducklings and eggs) in all three countries (Marinova-Petkova *et al*., 2016). Ducks often do not show clinical signs of disease, which makes identification of infection challenging. In addition, short production cycles, high movement of personnel and ducks between farms, and challenges in cleaning and disinfection of transport vehicles also contribute to potential maintenance of IAV without detection (Dent *et al*., 2011; Lowe *et al*., 2014; Marinova-Petkova *et al*., 2016; Guinat *et al*., 2019). Serological and swab testing of ducks as sentinel species might be useful in predicting future outbreaks in poultry.

Our analysis demonstrates movement of viruses in Bulgaria between farms, but the viruses are largely regionally sequestered. There is no evidence for movement between regions, for example, between Dobrich in the North-East and Plovdiv in Central Bulgaria. However, connections between Plovdiv and Eastern Bulgaria (Yambol, Sliven), which is geographically relatively close are clear. Vigilance should also be maintained for indirect transmission potential, as a more recently isolated virus from the western region of Vidin appears to be closely related to viruses found in Central Bulgaria, indicating a possible link between these regions either via wild birds or translocation on land.

Future studies with a focus on sampling domestic ducks and examining regional movement of people and commodities including equipment/vehicles between poultry farms within Bulgaria and trade links with poultry production systems in other European countries (especially Hungary and France) can help to understand the spread of HPAI viruses and design interventions.

## Supplementary Figures

**Supplementary Figure 1:**
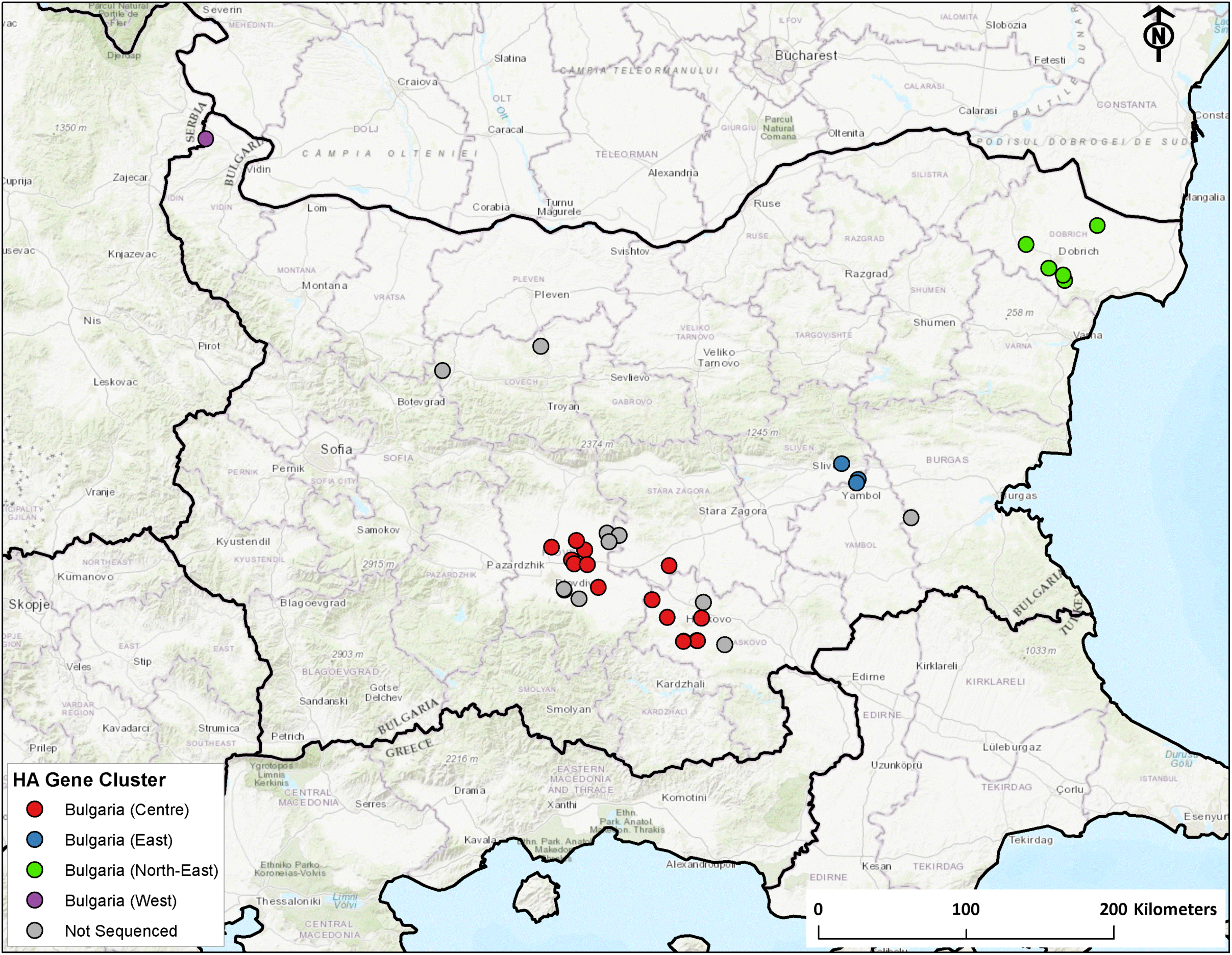
Map showing geographical distribution of viral genetic clusters in Bulgaria between 2017-2019

**Supplementary Figure 2:**
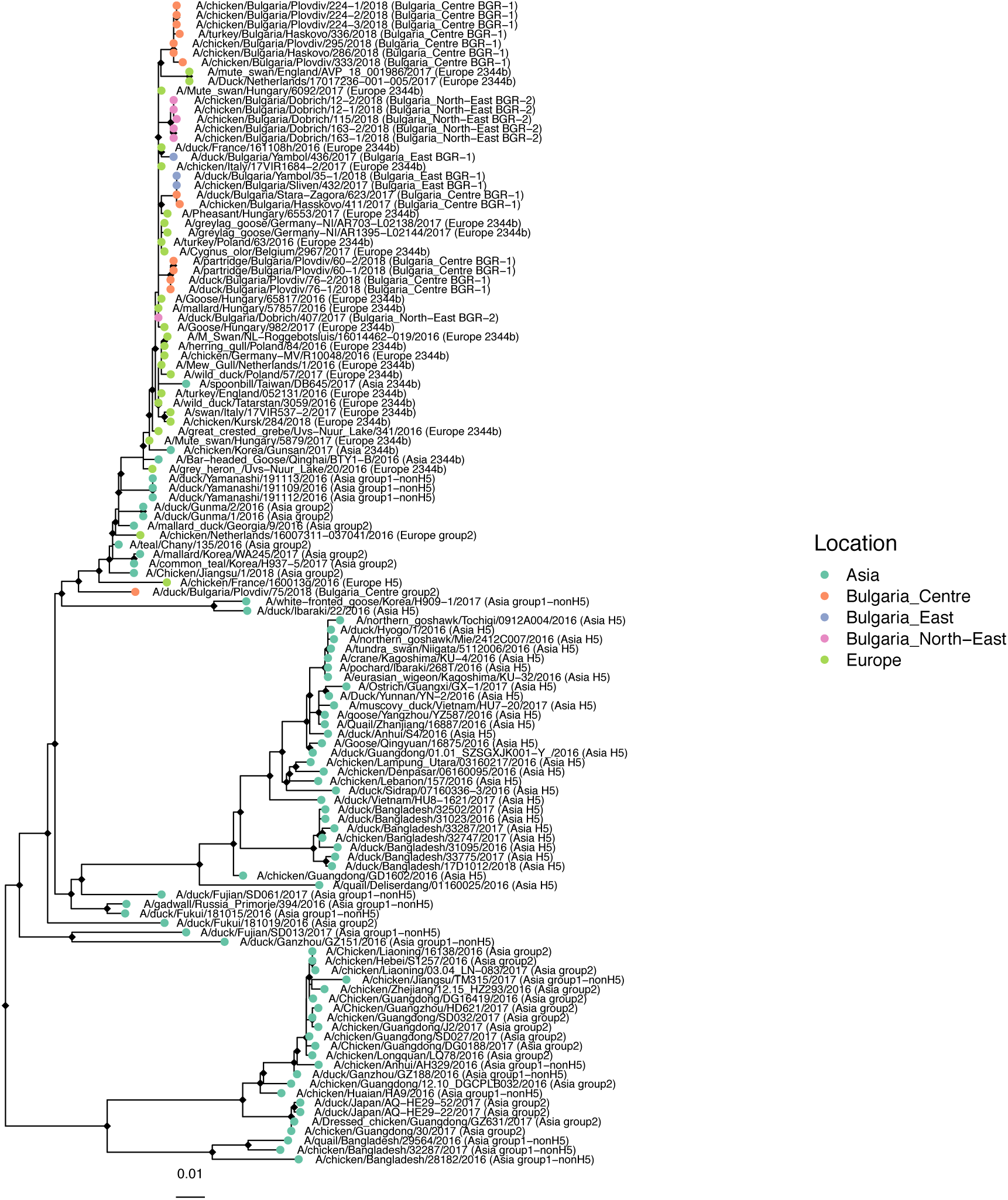
Maximum-likelihood tree for MP gene segment from Bulgarian (2017/18) and related viruses. Colours indicate location the virus was isolated from, and black diamonds at nodes indicate approximate likelihood ratio test (aLRT) branch supports > 0.9.

**Supplementary Table 1:** A record of highly pathogenic avian influenza disease events recorded in Bulgaria between 1 January 2019 and 8 April 2020. Table downloaded on 8 April 2020 from the FAO (Food and Agriculture Organisation) EMPRES-i (EMPRES Global Animal Disease Information System) initiative website: http://empres-i.fao.org/eipws3g/.

| Id | source | region | country | admin1 | locality Name | observation Date | reporting Date | status | disease | serotypes | species Description | sum AtRisk | sum Cases | sum Deaths | sum Destroyed |
| --- | --- | --- | --- | --- | --- | --- | --- | --- | --- | --- | --- | --- | --- | --- | --- |
| 266680 | EC | Europe | Bulgaria | Plovdiv | Trilistnik village | 12/03/2020 | 13/03/2020 | Confirmed | Influenza - Avian | H5N8 HPAI | domestic, chicken | 39120 | 50 |  |  |
| 266679 | EC | Europe | Bulgaria | Kardzhali | Kurdzhali municipal | 11/03/2020 | 13/03/2020 | Confirmed | Influenza - Avian | H5N8 HPAI | domestic, chicken | 16800 | 8000 |  |  |
| 266161 | OIE | Europe | Bulgaria | Plovdiv | Bulgaria | 02/03/2020 | 03/03/2020 | Confirmed | Influenza - Avian | H5N8 HPAI | domestic, unspecified bi | 5000 | 20 |  | 5000 |
| 266160 | OIE | Europe | Bulgaria | Plovdiv | Bulgaria | 02/03/2020 | 03/03/2020 | Confirmed | Influenza - Avian | H5N8 HPAI | domestic, unspecified bi | 3620 | 20 |  | 3620 |
| 265627 | OIE | Europe | Bulgaria | Plovdiv | Bulgaria | 24/02/2020 | 24/02/2020 | Confirmed | Influenza - Avian | H5N8 HPAI | domestic, duck | 11600 | 20 | 20 | 11580 |
| 265626 | OIE | Europe | Bulgaria | Plovdiv | Bulgaria | 24/02/2020 | 24/02/2020 | Confirmed | Influenza - Avian | H5N8 HPAI | domestic, chicken | 55437 | 34 | 34 | 55403 |
| 265136 | National authorities | Europe | Bulgaria | Plovdiv | Bulgaria | 17/02/2020 | 17/02/2020 | Confirmed | Influenza - Avian | H5N8 HPAI | domestic, duck | 15729 | 9142 | 9142 | 5830 |
| 250626 | National authorities | Europe | Bulgaria | Plovdiv | Asenovgrad | 08/04/2019 | 08/04/2019 | Confirmed | Influenza - Avian | H5N8 HPAI | domestic, chicken | 168752 | 300 | 300 | 168452 |
| 250532 | National authorities | Europe | Bulgaria | Plovdiv | Krumovo | 03/04/2019 | 05/04/2019 | Confirmed | Influenza - Avian | H5N8 HPAI | domestic, chicken | 37 | 37 | 37 | 0 |
| 250531 | National authorities | Europe | Bulgaria | Plovdiv | Asenovgrad | 03/04/2019 | 05/04/2019 | Confirmed | Influenza - Avian | H5N8 HPAI | domestic, chicken | 34998 | 298 | 298 | 34700 |
| 250202 | OIE | Europe | Bulgaria | Lovech | Lovech | 02/04/2019 | 03/04/2019 | Confirmed | Influenza - Avian | H5 HPAI | domestic, unspecified bi | 1400 | 20 | 0 | 1400 |
| 248811 | OIE | Europe | Bulgaria | Lovech | Lisets | 13/03/2019 | 13/03/2019 | Confirmed | Influenza - Avian | H5 HPAI | domestic, unspecified bi | 3200 | 20 | 0 |  |

